# Does circadian regulation lead to optimal gas exchange regulation?

**DOI:** 10.1101/121368

**Authors:** Víctor Resco de Dios, Arthur Gessler, Juan Pedro Ferrio, Josu G Alday, Michael Bahn, Jorge del Castillo, Sébastien Devidal, Sonia García-Muñoz, Zachary Kayler, Damien Landais, Paula Martín-Gómez, Alexandru Milcu, Clément Piel, Karin Pirhofer-Walzl, Olivier Ravel, Serajis Salekin, David T Tissue, Mark G Tjoelker, Jordi Voltas, Jacques Roy

## Abstract

Optimal stomatal theory is an evolutionary model proposing that leaves trade-off Carbon (C) for water to maximise C assimilation (*A*) and minimise transpiration (*E*), thereby generating a marginal water cost of carbon gain (λ) that remains constant over short temporal scales. The circadian clock is a molecular timer of metabolism that controls *A* and stomatal conductance (*g*_s_), amongst other processes, in a broad array of plant species. Here, we test whether circadian regulation contributes towards achieving optimal stomatal behaviour. We subjected bean (*Phaseolus vulgaris*) and cotton (*Gossypium hirsutum*) canopies to fixed, continuous environmental conditions of photosynthetically active radiation, temperature and vapour pressure deficit over 48 hours. We observed a significant and self-sustained circadian oscillation in *A* and in stomatal conductance (*g*_s_) which also led to a circadian oscillation in *λ*. The lack of constant marginal water cost indicates that circadian regulation does not directly lead to optimal stomatal behaviour. However, the temporal pattern in gas exchange, indicative of either maximizing *A* or of minimizing *E*, depending upon time of day, indicates that circadian regulation could contribute towards optimizing stomatal responses. More broadly, our results add to the emerging field of plant circadian ecology and show that molecular controls may partially explain leaf-level patterns observed in the field.

## Introduction

Early trade-offs in ecology recognized the need to balance growth and survival (Grubb 2015). Intense resource acquisition to sustain elevated growth rates, for instance, could lead to quick resource depletion and ultimately death. As a partial explanation for plant response to this constraint, the hypothesis of optimization in stomatal conductance was developed (Cowan 1977; Cowan & Farquhar 1977). In short, the optimal stomatal conductance hypothesis proposes that stomata balance the trade-off between *A* (C assimilation) and *E* (water losses) by maintaining a constant marginal water cost (*λ* = δ*E*/δ*A*; in mol CO_2_ mol^−1^ H_2_O), at least over short time scales, at the point where *A* is maximized and *E* minimized (Cowan & Farquhar 1977).

This optimal strategy was originally postulated as a conservative strategy for plants facing variation in a physical environment that, to a degree, is unpredictable or stochastic. As a result, most tests of the prediction of constant *λ* have been performed under changing vapour pressure deficit, soil water, temperature or [CO_2_] (Manzoni *et al.* 2011; Duursma *et al.* 2013; Buckley *et al.* 2014). However, there is also a degree of predictability in the variation observed in the natural environment. Chief amongst these is photoperiod, which varies deterministically as a function of day of year and of geographic location. A nearly universal adaptation to the photoperiod, and other predictable environmental cues, is the endogenous circadian clock (McClung 2006; Resco, Hartwell & Hall 2009).

Circadian rhythms regulate the transcription of ∼30% of the plant’s genome (Covington *et al.* 2008) and, amongst others, diurnal patterns of stomatal conductance and photosynthesis are partially products of circadian regulation (Hennessey, Freeden & Field 1993; Mencuccini, Mambelli & Comstock 2000). It has been shown that resonance between circadian rhythms in gas exchange and environmental cues increases plant growth (Graf *et al.* 2010; de Montaigu *et al.* 2015; Kolling *et al.* 2015; Resco de Dios *et al.* 2016), and that circadian timing is related to photosynthesis rates and stomatal conductance (Edwards *et al.* 2011). Regarding the hypothesis of optimal stomatal regulation, Cowan (1982) states that “if diurnal variation in natural physical environment were regular and predictable, then optimization would require only that there be an appropriate circadian rhythm in stomatal aperture”. Given that variation in the physical environment is not entirely regular and predictable, here we seek to understand the potential role of circadian rhythms towards optimizing the trade-off in *A* vs *E*.

Circadian biologists often mention circadian regulation as an important component of achieving optimal stomatal conductance (Hubbard & Webb 2015). However, we are unaware of any direct tests for optimality resulting from circadian regulation, and perhaps the word optimal in those studies is used in general terms, and not in relation to the specific hypothesis of time-invariant *λ*. In fact, circadian regulation in *A* has been documented to be uncoupled and independent from circadian regulation in *g*_s_ (Dodd, Parkinson & Webb 2004), but linkages between these two processes is a pre-requisite for optimal water use. Therefore, if circadian rhythms regulate *A* and *g*_s_ independently from each other, one would hypothesize that circadian regulation alone, would not lead to optimal stomatal regulation.

Nonetheless, there is some evidence from theoretical modeling that circadian rhythms could aid in reaching optimality. Circadian regulation serves to “anticipate” predictable environmental cues, in such a way that stomata can adjust in advance (“stomatal priming”, Resco de Dios *et al.* 2016). As such, the clock has been hypothesized to aid in attaining optimality through stomatal priming because direct responses to regular diurnal fluctuations alone would inevitably lead to a lagged response (Dietze 2014). In other words, stomata show a lagged response to the environment (Vico *et al.* 2011) and, although it is not expected that optimality operates at every instant, circadian regulation could help in achieving optimality by diminishing the lags through stomatal priming (Dietze 2014).

Here, we propose that circadian regulation, *per se*, does not lead to optimal behavior over diurnal cycles, but that it might help in achieving optimality within field settings. More explicitly, we hypothesize that: 1) because the circadian clock regulates *A* and *g*_s_ independently, circadian action will lead to a time-changing *λ*, consistent with non-optimal behavior; 2) the temporal pattern of circadian driven gas exchange will be consistent with a stomatal priming that prepares for regular environmental variation.

Assessing the effects of circadian regulation on daytime *A* and *g*_s_ under natural conditions is difficult because the influence of environmental drivers generally mask circadian regulation. Circadian regulation is most strongly expressed under a “constant environment”: when temperature, radiation, vapour pressure deficit and other environmental drivers are held experimentally constant over 24h or longer. Therefore, we addressed our questions by examining temporal variation in *λ* in an herb (bean, *Phaseolus vulgaris*) and in a shrub (cotton, *Gossypium hirsutum*) under 48h of constant environmental conditions.

It has been noted that the optimal stomatal hypothesis cannot be tested directly with experimental manipulations such as 48h of constant environmental conditions, because optimal stomatal theory represents an evolutionary process and therefore can only be assessed under environmental conditions observed during their evolution (Cowan 2002). However, tests of the optimal stomatal hypothesis have been successfully conducted in elevated CO_2_ enrichment experiments (Barton *et al.* 2012; Medlyn *et al.* 2013), although it is clear that plants have not evolved experiencing step-function, large sudden increases in CO_2_ concentration (Woodward 2007). We suggest that our experimental approach is somewhat similar to other approaches, except that to avoid the potential for experimental artefacts, we do not test whether the optimal stomatal theory is observed under constant environmental conditions. Instead, our goal is to assess the potential for circadian regulation to contribute to optimal stomatal behaviour in natural, varying environments.

Throughout the manuscript, we will present data on both the marginal water cost of carbon gain (*λ* = δ*E*/δ*A*) and on *A*_net_/*g*_s_ (intrinsic water use efficiency). This is for the sake of clarity as *A*_net_/*g*_s_ is more often used than the marginal water cost of carbon gain, and also because those two variables tend to be inversely correlated (using the convention of this manuscript, see methods for calculation of *λ*). However, it is important to note that *λ* is not simply the inverse of water use efficiency (*A*/*E*) because *λ* is a partial derivative, i.e. an expression of the co-variation between the two variables for a given level of *g*_s_ (see for instance (Thomas, Eamus & Bell 1999) for different methods of calculation).

## Materials and methods

### Experimental set-up

The experiment was performed at the Macrocosms platform of the Montpellier European Ecotron, Centre National de la Recherche Scientifique (CNRS, France). We used 6 controlled-environment units of the macrocosms platform (3 planted with bean and 3 with cotton), where the main abiotic (air temperature, humidity and CO_2_ concentration) drivers were automatically controlled. The soil was extracted using large lysimeters (2 m^2^, circular with a diameter of 1.6 m, weighing 7 to 8 tonnes) from the flood plain of the Saale River near Jena, Germany, and used in a previous Ecotron experiment on biodiversity (Milcu *et al.* 2014). After that experiment, the soil was ploughed down to 40 cm and fertilized with 25/25/35 NPK (MgO, SO_3_ and other oligoelements were associated in this fertilizer: Engrais bleu universel, BINOR, Fleury-les-Aubrais, FR).

The soil was regularly watered to *ca.* field capacity by drip irrigation, although irrigation was stopped during each measurement campaign (few days) to avoid interference with water flux measurements. However, no significant differences (at *P* < 0.05, paired t-test, n = 3) in leaf water potential occurred between the beginning and end of these measurement campaigns, indicating no apparent effect of a potentially declining soil moisture on leaf hydration.

Environmental conditions within the macrocosms (excluding the experimental periods) were set to mimic outdoor conditions, but did include a 10% light reduction by the macrocosm dome cover (sheet of Fluorinated Ethylene Propylene). During experimental periods, light was controlled by placing a completely opaque fitted cover on each dome to block external light inputs (PVC coated polyester sheet Ferrari 502, assembled by IASO, Lleida, Spain), and by using a set of 5 dimmable plasma lamps (GAN 300 LEP with the Luxim STA 41.02 bulb, with a sun-like light spectrum); these lamps were hung 30 cm above the plant canopy and provided a PAR of 500 µmol m^−2^ s^−1^. We measured PAR at canopy level with a quantum sensor (Li-190, LI-COR Biosciences, Lincoln, NE, USA) in each macrocosm.

Bean and cotton were planted in 5 different rows within the macrocosms on 10^th^ July 2013, one month before the start of the measurements, and thinned to densities of 10.5 and 9 individuals m^−2^, respectively. Cotton (STAM-A16 variety by INRAB/CIRAD) is a perennial shrub with an indeterminate growth habit. This cotton variety grows to 1.5-2 m tall and has a pyramidal shape and short branches. Bean (recombinant inbred line RIL-115 bred by INRA Eco&Sol) is an annual herbaceous species. RIL-115 is a fast growing, indeterminate dwarf variety, 0.3-0.5 m tall; it was inoculated with *Rhizobium tropici* CIAT 899 also provided by INRA. During the experiment, bean and cotton generally remained at the inflorescence emergence developmental growth stage (codes 51-59 in BBCH scale, the standard phenological scale within the crop industry; Feller *et al.* 1995; Munger *et al.* 1998). Further details on Ecotron measurements have been provided elsewhere (Resco de Dios *et al.* 2015).

During each experimental period, plants were entrained for five days under environmental conditions that mimicked the pattern observed in an average August sunny day in Montpellier in terms of *T*_air_ (28/19° C, diurnal max/min) and VPD. However, we kept radiation levels much lower (at 500 µmol m^−2^ s^−1^ at canopy level) because previous research proposed that stomatal behavior should follow optimal theory when photosynthesis is light- (and not CO_2_) limited (Medlyn *et al.* 2011), and our PAR values ensured this was the case. After 5 days of entrainment, we maintained environmental conditions constant starting at solar noon and for the next 48 h.

### Measurements

We characterized the general pattern of gas exchange during the last day of entrainment and during constant environmental conditions by monitoring, every 4 hours, gas exchange (LI-6400XT, Li-Cor, Lincoln, Nebraska, USA) in three different leaves, within each of the three domes, per species that were available. To diminish redundancy in the presentation, only *A*_net_/*g*_s_ will be shown to characterize this general pattern.

To obtain enough resolution to test for temporal changes in the marginal water cost of carbon gain, we additionally measured gas exchange every 2 minutes by using 2-3 additional portable photosynthesis systems per species and day. Each instrument was continuously deployed on a leaf for 24 h, and the Auto-Log function was used. Measurements were conducted over 48 h with an effective *n* = 3 per species (1-2 leaves were measured per macrocosm, in a total of 3 macrocosms). To diminish redundancy in the presentation, only *A*_net_ and *g*_s_ (along with *g*_1_) will be shown from these high-resolution measurements (but not *A*_net_/*g*_s_).

### Analyses

The marginal water cost used was estimated from parameter *g*_1_ in the stomatal model of Medlyn *et al.* (2011). This model was derived from optimal stomatal theory, such that the *g*_1_ is inversely proportional to the root square of *λ*. We calculated the marginal water cost from the Medlyn *et al.* (2011), and assuming minimal conductance (*g*_0_) was 0, so that we could compare the variability observed in our experiment with that observed in a recent synthesis reporting *g*_1_ values for 314 species from 10 different functional types (Lin *et al.* 2015). We calculated values of *g*_1_ separately for each hour.

We examined statistical significance of temporal patterns in *g*_1_ with a Generalized Additive Mixed Model (GAMM) fitted with automated smoothness selection (Wood 2006) in the R software environment (*mgcv* library in R 3.1.2, The R Foundation for Statistical Computing, Vienna, Austria), including macrocosms as a random factor. This approach was chosen because there are no *a priori* assumptions about the functional relationship between variables. We accounted for temporal autocorrelation in the residuals by adding a first-order autoregressive process structure (nlme library, Pinheiro & Bates 2000). Significant temporal variation in the GAMM best-fit line was analysed after computation of the first derivative (the slope, or rate of change) with the finite differences method. We also computed standard errors and a 95% point-wise confidence interval for the first derivative. The trend was subsequently deemed significant when the derivative confidence interval was bounded away from zero at the 95% level (for full details on this method see Curtis & Simpson 2014). Periods with no significant variation are illustrated on the figures by the yellow line portions, and significant differences occur elsewhere. The magnitude of the range in variation driven by the circadian clock was calculated using GAMM maximum and minimum predicted values.

## Results

We observed a self-sustained oscillation in *A*_net_, *g*_s_ and *A*_net_/*g*_s_ that showed a ∼24 h periodicity (Figs. 1, 2, Table 1). That is, there was a significant variation in *A*_net_ and *g*_s_ in the absence of variation in environmental drivers, and this variation showed a diurnal cycle. Although *A*_net_ and *g*_s_ generally followed the same pattern in that they both concurrently showed either a positive or a negative trend, the magnitude of the oscillation was larger in *g*_s_ (54-84% change, Table 1), than in *A*_net_ (28-42% change, Table 1) over a 24 h cycle in constant environmental conditions. In turn, this led to a significant variation in instantaneous water use efficiency (*A*_net_/*g*_s_) that was 46-74% of that during entrainment (Table 1). If we only consider the oscillation during the subjective day (the time under constant conditions when it would have normally been daytime during entrainment) we still observe a significant and time-dependent variation in *A*_net_, *g*_s_ and *A*_net_/*g*_s_, although of smaller magnitude than during the whole 24 h cycle (Figs. 1, 2).

**Figure 1:**
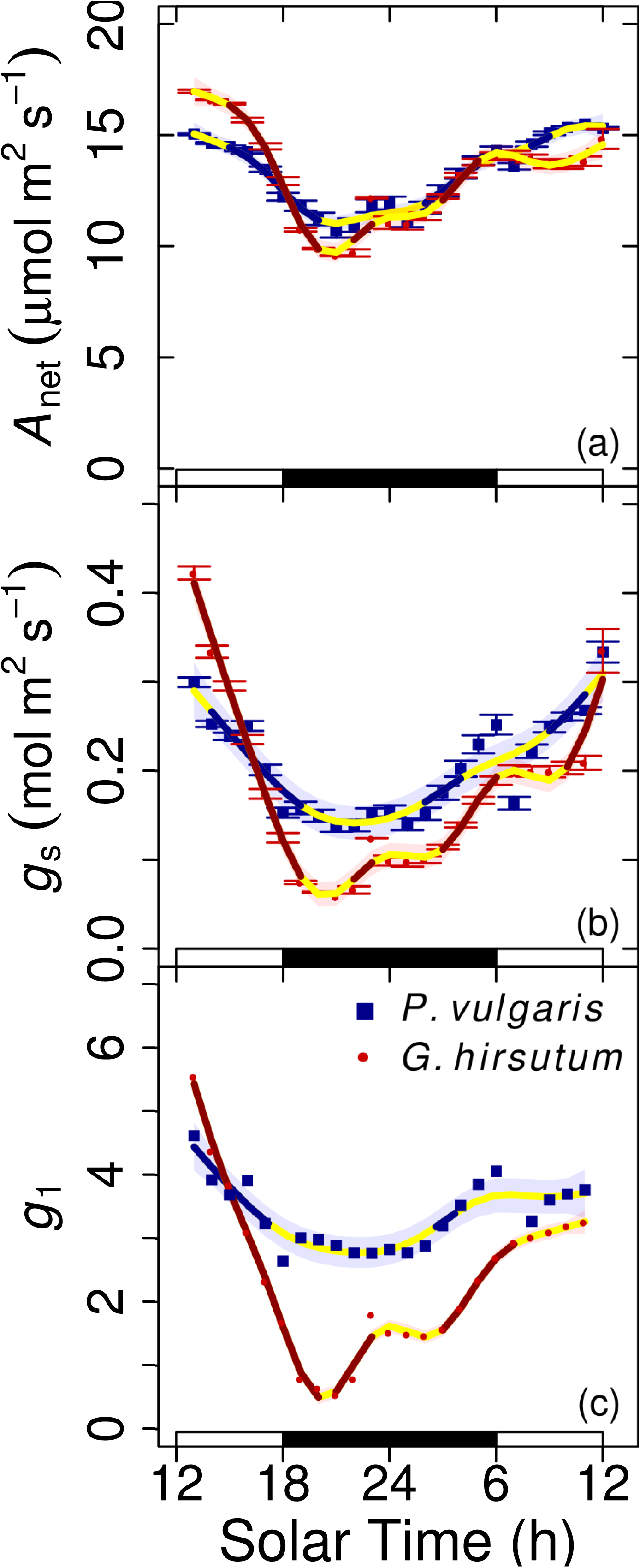
Circadian oscillation in the marginal water cost. The dots (with small SE bars hidden) indicate hourly values of assimilation (*A*_net_) stomatal conductance (*g*_s_) and a parameter proportional to the marginal water cost of carbon gain (*g*_1_). Measurements were taken concomitantly to those under constant conditions reported in Fig. 2, although data from both days were pooled together to increase sample size. The white and black rectangles at the base indicate the subjective day (when it would have been daytime during entrainment) and subjective night, respectively, under constant conditions. Lines (and shaded error intervals) indicate the prediction (and SE) of Generalized Additive Model (GAM) fitting separately for each species (some lines may overlap), and portions which are not yellow indicate significant temporal variation.

**Figure 2.**
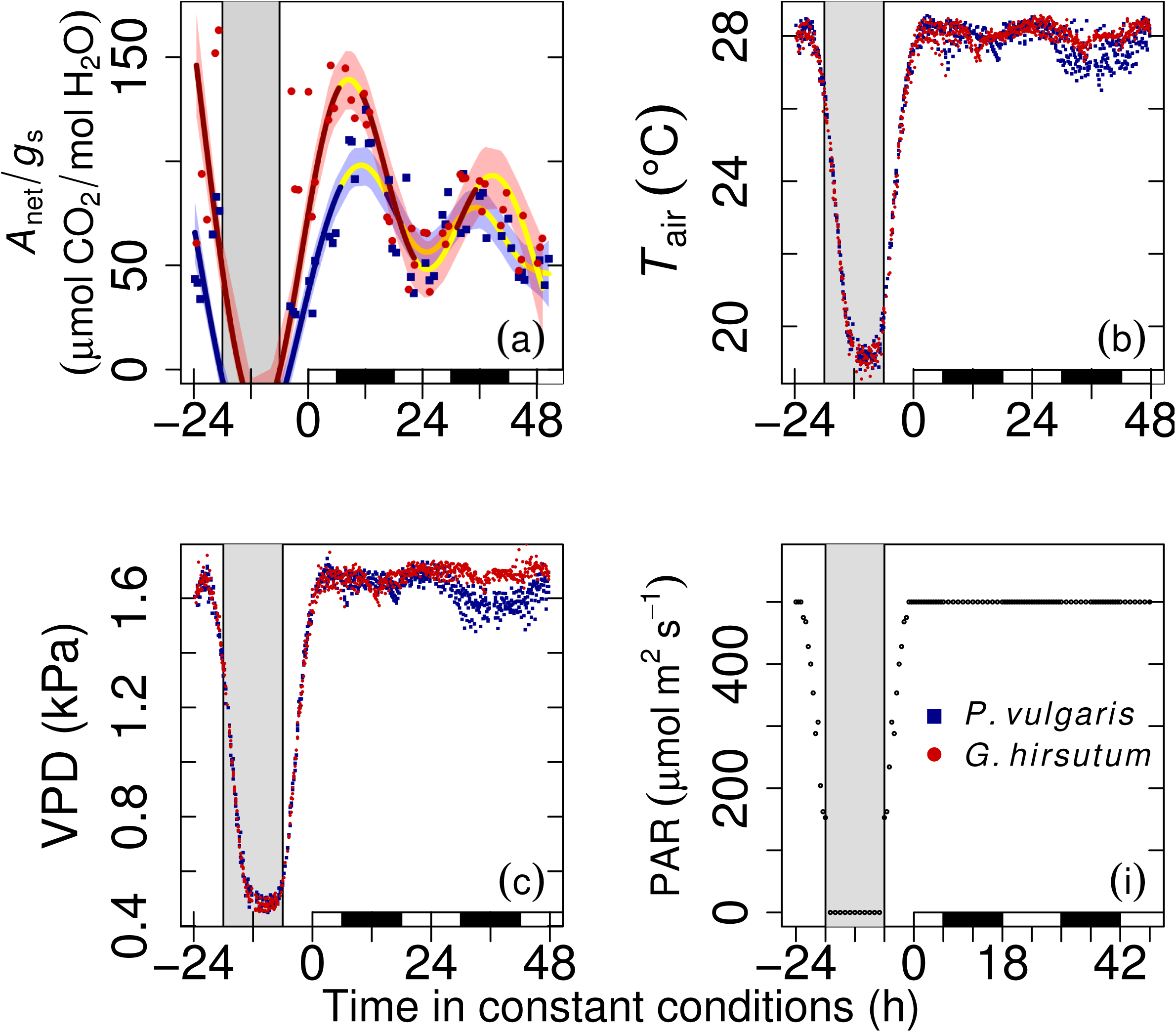
Circadian regulation of leaf assimilation (*A*_net_) over stomatal conductance (*g*_s_). During entrainment, environmental conditions of Temperature (*T*_air_) and Vapor Pressure Deficit (VPD) mimicked those outdoors, with 500 µmol m^−2^ s^−1^ PAR (first 24 h shown), and remained constant for 48 h starting at solar noon. The grey and white backgrounds indicates when PAR was at 0 (µmol m^−2^ s^−1^) or higher, respectively. White and black rectangles at the base indicate the subjective day (when it would have been daytime during entrainment) and subjective night, respectively, under constant conditions. Dots represent measured values at each of three replicate macrocosms, and thick lines (and shaded error intervals) indicate the prediction (and SE) of Generalized Additive Mixed Model (GAMM) fitting separately for each species (some lines may overlap). GAMM best-fit line portions which are not yellow indicate significant temporal variation. Values in (b-d) were measured by the different ms every 15 minutes and values in (a) were measured every 4 hours with a hotosynthesis system (the cuvette was set to match macrocosm conditions).

**Table 1:** Quantification of the circadian-driven range in variation of diurnal gas exchange. The variation in fluxes under constant environmental conditions was derived from Generalized Aditive Mixed Model predictions in Fig. 2.

| Process | Species | Variation under constant conditions |  |  |
| --- | --- | --- | --- | --- |
|  |  | Maximum | Minimum | % Change |
| $A_{\text{net}}$ ($\mu\text{mol m}^{-2} \text{s}^{-1}$) | *P. vulgaris* | 15.42 | 11.02 | 28.48 |
|  | *G. hirsutum* | 16.97 | 9.73 | 42.65 |
| $g_s$ ($\text{mol m}^{-2} \text{s}^{-1}$) | *P. vulgaris* | 0.31 | 0.14 | 54.03 |
|  | *G. hirsutum* | 0.41 | 0.063 | 84.85 |
| $A_{\text{net}}/g_s$ ($\mu\text{mol CO}_2 / \text{mol H}_2\text{O}$) | *P. vulgaris* | 95.37 | 51.17 | 46.35 |
|  | *G. hirsutum* | 156.23 | 40.52 | 74.06 |

The pattern in *A*_net_/*g*_s_ was such that water use efficiency increased in the first subjective afternoon under constant conditions (hours 0-6 in Fig. 2c), it remained constant in the first hours of the night, and then it increased again from the subjective midnight (hour 12 in Fig. 2c) until the following subjective noon (hour 24 in Fig. 2c). The rhythm dampens slightly in the second 24 h period under constant conditions (hours 24-48), because the clock is flexible and becomes entrained every day (Hennessey & Field 1991; Graf *et al.* 2010). However, we can still observe temporal fluctuations similar to those in the previous day (although with a smaller degree of significance). Because this study mostly focused on the implications of clock regulation within field settings, the results during the first day are most important because this reflects the period of highest influence of natural environmental variation.

We also observed a diurnal self-sustained cycle in *g*_1_ (Fig 1). That is, we did not observe homeostasis in the marginal water cost despite lack of variation in environmental drivers. Instead, we observed a pattern that was generally opposite to that found in water use efficiency: *g*_1_ significantly declined during the subjective afternoon in both species (although with a more pronounced decline in cotton), and a significant increase during the subjective night occurred for both species, that continued into the subjective morning for cotton.

## Discussion

### The importance of circadian regulation towards achieving optimization

We observed a significant and self-sustained 24 h oscillation in *A*_net_ and *g*_s_, of different magnitude for each process, and that ultimately led to a diurnal oscillation in intrinsic water use efficiency and in the marginal cost of water, despite the absence of variation in environmental drivers. Diurnal variation in *g*_1_ ranged from 5.5 to 0.5 over the 24 h cycle, and from 5.5 to 1.7 when we only consider variation during the subjective daytime (Fig. 1). There are many processes that could explain an afternoon decline in *A*_net_, including feedback inhibition from starch accumulation, photorespiration and stomatal feedbacks, amongst others (Azcón-Bieto 1983; Jones 1998; Flexas *et al.* 2006). Similarly, a multitude of processes could explain the afternoon decline in *g*_s_, including hydraulic feedbacks and depletion of stem capacitors (Jones 1998; Zhang *et al.* 2014). However, the only process that can explain a self-sustained 24 h cycle is the circadian clock (Resco, Hartwell & Hall 2009). We can therefore conclude that, in the absence of variation in the physical environment, circadian regulation of stomatal behaviour *per se* does not directly lead to an optimization, because *g*_1_ was not constant throughout the experiment. However, as we will discuss further below, the pattern of variation in *g*_1_ indicates that circadian regulation could be an important contributor to achieving optimality in the field.

Under the well-watered and fertilized conditions of this experiment, where radiation was probably the only limiting factor, we observed a stronger relative fluctuation in *g*_s_ than in *A*_net_. This result is consistent with previous studies (Doughty *et al.* 2006; Yakir *et al.* 2007) and suggests that stronger clock regulation over *g*_s_ than over *A*_net_ could be widespread across C_3_ plants. On the one hand, these temporal patterns could be interpreted as an indication that the clock fosters a maximization of *A* at the time of maximal potential for assimilation (*A* peaked at the subjective noon) which, in turn, would be aided by a maximal *g*_s_ which decreases diffusional limitations. On the other hand, the stronger decrease in *g*_s_, relative to that in *A*_net_, during the subjective morning and afternoon, when conditions would have become less favourable for assimilation in a naturally fluctuating environment, is consistent with a conservative water use strategy. Therefore, this result is consistent with the hypothesis that circadian-driven stomatal priming could contribute towards reaching optimality (Dietze 2014).

Nonetheless, this stomatal regulation strategy contrasts with other work that has shown that circadian regulation tends towards “wasting” water at times when there is no *A*. Circadian regulation is one of the main drivers of the temporal pattern of nocturnal *g*_s_, which increases constantly from *ca.* midnight until predawn (Caird, Richards & Donovan 2007; Resco de Dios et al. 2015). There is no *A* overnight and, therefore nocturnal water use is not directly linked with assimilation. However, different studies have linked higher predawn *g*_s_ with higher *A* early in the morning (although whether or not this has a significant effect on plant growth is still under discussion, cf. Auchincloss *et al.* 2014; Resco de Dios *et al.* 2016). Therefore, compensation could occur if we observe circadian regulation over the full diurnal cycle, where an increase in *A* at different times (*e.g*: subjective noon in this study or early morning in the cited work) would be accompanied by higher water losses (maximum *g*_s_ at subjective noon, and increase in *g*_s_ overnight, respectively); but a more conservative water use occurs at other times (when the relative decline in *g*_s_ is higher than in *A*, such as the afternoon or evening).

Our study is, to the best of our knowledge, the first to report a circadian pattern in *g*_1_. As previously mentioned, the optimal stomatal hypothesis does not present specific predictions about what should happen under environmental conditions that do not naturally occur in the field (Cowan 2002). One could argue that 24h of continuous light does occur above the Polar circle, but not as a constant light intensity as utilised in our experiment, and moreover, bean and cotton did not evolve in these constant light environments. However, all species experience cloudy days over their lifespans. Under cloudy days, temporal variation in temperature, vapour pressure deficit and other environmental drivers is generally minimal. Therefore, plants do often experience environmental conditions that are roughly constant for a few hours. It is therefore notable that the largest change in *g*_1_ occurred in the first 6 hours after conditions were kept constant (1200h to 1800h solar time) and this change in *g*_1_ (from 5.5 to 1.7, see above) was significant. In fact, a recent global synthesis shows that mean *g*_1_ values across different functional types (in a study encompassing 314 species) ranged between 1.6 and 7.2 (Lin *et al.* 2015). Subsequently, we encourage field studies of leaf-level gas exchange conducted at high temporal resolution to assess the extent of temporal variation in *g*_1_ under cloudy days.

### Implications and mechanisms

It has been argued that more biological realism must be incorporated into optimality models to generate a better understanding of optimal behaviour and its constraints (Niinemets 2012). Our results indicate that circadian regulation might be one of the most important processes to be included in these models. For instance, it is well documented that hysteresis in the *E*-VPD relationship generally exists, with higher *E* values in the morning than in the afternoon, at any given VPD. There are different processes that could explain this phenomenon (O’Grady, Eamus & Hutley 1999; Tuzet, Perrier & Leuning 2003; Unsworth *et al.* 2004) and, one of them, is the lag between peaks in radiation and VPD (radiation peaks at solar noon, but VPD peaks a few hours later) (Zhang *et al.* 2014). Circadian rhythms could contribute to this phenomenon. The clock is often considered to be entrained by both temperature and radiation (Millar 2004). However, the pattern of *A* and *g*_s_ resembles more closely that of radiation, in that both *A* and *g*_s_ peaked at subjective noon, which was the same time for PAR during entrainment. However, *T*_air_ and VPD peaked at 1400h during entrainment, and circadian regulation would have already started to decrease stomatal conductance at that time. Therefore, circadian-driven stomatal closure after radiation peaks at noon (which are more pronounced than the decline in *A*), in concert with radiation-VPD lags, could be a contributing factor in the documented hysteresis in E-VPD relationships; however, this is not currently accounted for in models.

Circadian clocks in plants have traditionally been assumed to be cell autonomous and not coordinated across cells or plant tissues (Endo *et al.* 2014). However, recent research has observed that a hierarchy exists in plants in that the clock in the leaf vascular tissue regulates the clock in the mesophyll leaf tissue (Endo *et al.* 2014). Although speculative, it is tempting to hypothesize that clock-controlled hydraulic signals over vascular tissue could also be part of the response driving hysteresis in diurnal transpiration cycles.

The effect of circadian regulation on stomatal physiology is still being debated. In *Arabidopsis*, it has been proposed that the central oscillator of the clock directly controls stomatal behaviour because TOC1 (a component of the central oscillator) regulates ABA signalling (Legnaioli, Cuevas & Mas 2009). However, other studies have documented that time-dependent circadian regulation of *g*_s_ is independent of ABA concentration in beans (Mencuccini, Mambelli & Comstock 2000). Another line of research proposes that it is through [Ca_2+_]_cyt_ signalling that the circadian clock regulates stomatal movements (Hubbard & Webb 2015). Circadian regulation of *A* is relatively better understood, and it involves the joint regulation of the light harvesting complex, the carboxylating enzyme Rubisco, and feedbacks from carbohydrates (Dodd *et al.* 2014). However, most studies have been conducted at the molecular level with *Arabidopsis*, and the mechanism of action at “phenotypic” or eco-physiological scales, as well as the degree to which processes in *Arabidopsis* are generalizable to other species, remain unknown.

### Conclusions

It has been known for long that the circadian clock could be an important an important component underlying plant fitness. Understanding the reason why the circadian clock is adaptive has proven more challenging. Here we developed the first formal test of the hypothesis that the circadian clock leads towards optimal stomatal regulation and, indeed, the strong stomatal regulation under constant environmental conditions points to the circadian clock as an important component. Although we did not observe a constant marginal water cost under constant conditions, which is necessary for stomatal regulation to be optimal, the optimal stomatal hypothesis would also not have predicted that to occur given the artificiality of the experimental treatment. Importantly, the temporal patterns observed indicate how variation in stomatal regulation was consistent with a circadian-driven stomatal priming that prepares gas exchange in advance of regular environmental fluctuation. Although our experiments were not conducted under conditions typical of field settings, the strong fluctuation in *A* and *g*_s_ indicate that circadian regulation could be an important component underlying optimal behaviour in the field. These results add to the emerging field of plant circadian ecology and show that one of the mechanisms by which the circadian clock increases plant fitness is by contributing towards reaching optimal stomatal behaviour. Further studies will need to clarify whether the large changes observed in *g*_1_ under the subjective afternoon also occur in other species and under cloudy conditions.

## Acknowledgements

This study benefited from the CNRS human and technical resources allocated to the ECOTRONS Research Infrastructures as well as from the state allocation ‘Investissement d’Avenir’ AnaEE-France ANR-11-INBS-0001, ExpeER Transnational Access program, Ramón y Cajal fellowships (RYC-2012-10970 to VRD and RYC-2008-02050 to JPF) and internal grants from UWS-HIE to VRD and ZALF to AG. We remain indebted to E. Gerardeau, D. Dessauw, J. Jean, P. Prudent (Aïda CIRAD), C. Pernot (Eco&Sol INRA), B. Buatois, A. Rocheteau (CEFE CNRS), A. Pra, A. Mokhtar and the full Ecotron team, in particular C. Escape, for outstanding technical assistance during experiment set-up, plant cultivation or subsequent measurements. We also acknowledge R. Duursma, B. Medlyn for comments on an earlier version of this manuscript and to Y.-S. Lin and G.D. Farquhar for useful discussion.

## Data Accessibility

Data are freely accessible upon registration from http://www.ecotron.cnrs.fr/index.php/en/component/users/?view=login.

